# HSV-1 ICP0 Dimer Domain Adopts a Novel β-barrel Fold

**DOI:** 10.1101/2024.01.16.575752

**Authors:** Erick McCloskey, Maithri Kashipathy, Anne Cooper, Philip Gao, David K. Johnson, Kevin P. Battaile, Scott Lovell, David J. Davido

**Affiliations:** Department of Molecular Biosciences, University of Kansas, Lawrence, KS, USA; Protein Structure and X-Ray Crystallography Laboratory, University of Kansas, Lawrence, KS, USA; Protein Production Group, University of Kansas, Lawrence, KS, USA; Chemical Computational Biology Core, University of Kansas, Lawrence, KS, USA; NYX, New York Structural Biology Center, Upton, NY 10027, USA

**Author notes:** **CORRESPONDING AUTHOR INFORMATION** David Davido.

**Keywords:** HSV-1, ICP0, beta-barrel, crystal structure, novel fold, dimerization

## Abstract

Infected cell protein 0 (ICP0) is an immediate-early regulatory protein of herpes simplex virus 1 (HSV-1) that possesses E3 ubiquitin ligase activity. ICP0 transactivates viral genes, in part, through its C-terminal dimer domain (residues 555-767). Deletion of this dimer domain results in reduced viral gene expression, lytic infection, and reactivation from latency. Since ICP0’s dimer domain is associated with its transactivation activity and efficient viral replication, we wanted to determine the structure of this specific domain. The C-terminus of ICP0 was purified from bacteria and analyzed by X-ray crystallography to solve its structure. Each subunit or monomer in the ICP0 dimer is composed of nine β-strands and two α-helices. Interestingly, two adjacent β-strands from one monomer “reach” into the adjacent subunit during dimer formation, generating two β-barrel-like structures. Additionally, crystallographic analyses indicate a tetramer structure is formed from two β-strands of each dimer, creating a “stacking” of the β-barrels. The structural protein database searches indicate the fold or structure adopted by the ICP0 dimer is novel. The dimer is held together by an extensive network of hydrogen bonds. Computational analyses reveal that ICP0 can either form a dimer or bind to SUMO1 via its C-terminal SUMO-interacting motifs but not both. Understanding the structure of the dimer domain will provide insights into the activities of ICP0 and, ultimately, the HSV-1 life cycle.

## INTRODUCTION

It is estimated that 70%^1^ of the world’s population is infected with herpes simplex virus 1 (HSV-1), and infections in humans can cause painful oral and genital sores^2^. Additionally, HSV-1 is the major cause of spontaneous viral encephalitis^2^. While rare, this can be life-threatening, especially in newborns^2^. HSV-1 is the primary cause of infectious blindness in western industrialized countries, as infection of the cornea can lead to herpetic stromal keratitis (HSK)^2^. HSK and some cases of encephalitis can result from reactivation of an HSV-1 latent infection. In order to establish a latent infection, HSV-1 must first infect the nervous system from the periphery to subsequently reactivate and cause recurrent disease.

One of five immediate-early (IE) viral regulatory proteins that plays an important role in lytic and latent infections is infected cell protein 0 (ICP0). ICP0 is a 776-amino acid multi-functional protein that acts as a potent transactivator of viral gene expression^3–7^. ICP0 viral mutants grow poorly in cell culture, in animal models of HSV infection, and are impaired for reactivation^8–12^. Transient transfection assays and genetic studies have shown that the C-terminus of ICP0, which contains its C-terminal dimer domain (residues 555-767)^13^, contributes to its transactivator function and productive replication in cell culture^14,^ and in a mouse model of HSV-1 infection^15^. The mechanism of how ICP0 dimerization regulates viral gene expression has not yet been established.

The C-terminal domain (CTD) of ICP0 is multi-functional. **Figure 1A** shows the primary structure of ICP0 with several known domains or motifs. The RING-finger domain near the N-terminus is included for reference. The nuclear localization signal (NLS) is a short sequence of residues, 501-507. Ubiquitin specific protease 7 (USP7), which has the ability to remove conjugated ubiquitin from target proteins, binds in the domain from residues 619-635, an interaction that is associated with ICP0 stability^16,17^. The CoREST binding domain is defined from residues 667-739. CoREST is a component of the histone deacetylase 1/2 (HDAC1/2)–CoREST–REST repressor complex, shown to be disassociated by interacting with ICP0 and has been proposed to counteract viral gene silencing^18^. The minimal domain required for dimerization has been reported to be from amino acids 618-712^13^. The entire ICP0 dimerization domain is from residues 555-767, and it overlaps with three SUMO (small ubiquitin-like modifier)-binding sites, referred to as SUMO-interacting motif (SIM)-like sequences (SLS): SLS5, 6, and 7^19^. The nuclear domain 10 (ND10) localization region (residues 634-768) overlaps these SUMO-binding sequences^20^. ND10 bodies are part of intrinsic host defenses, as they contain several cellular proteins that play roles in protein cell cycle, stress, and interferon signaling, and are associated with repressing HSV-1 transcription^19,21^. ICP0’s disruption of ND10 organelles in the nucleus is, in part, by targeting SUMO or SUMO-modified proteins^19,22,23^.

**Figure 1:**
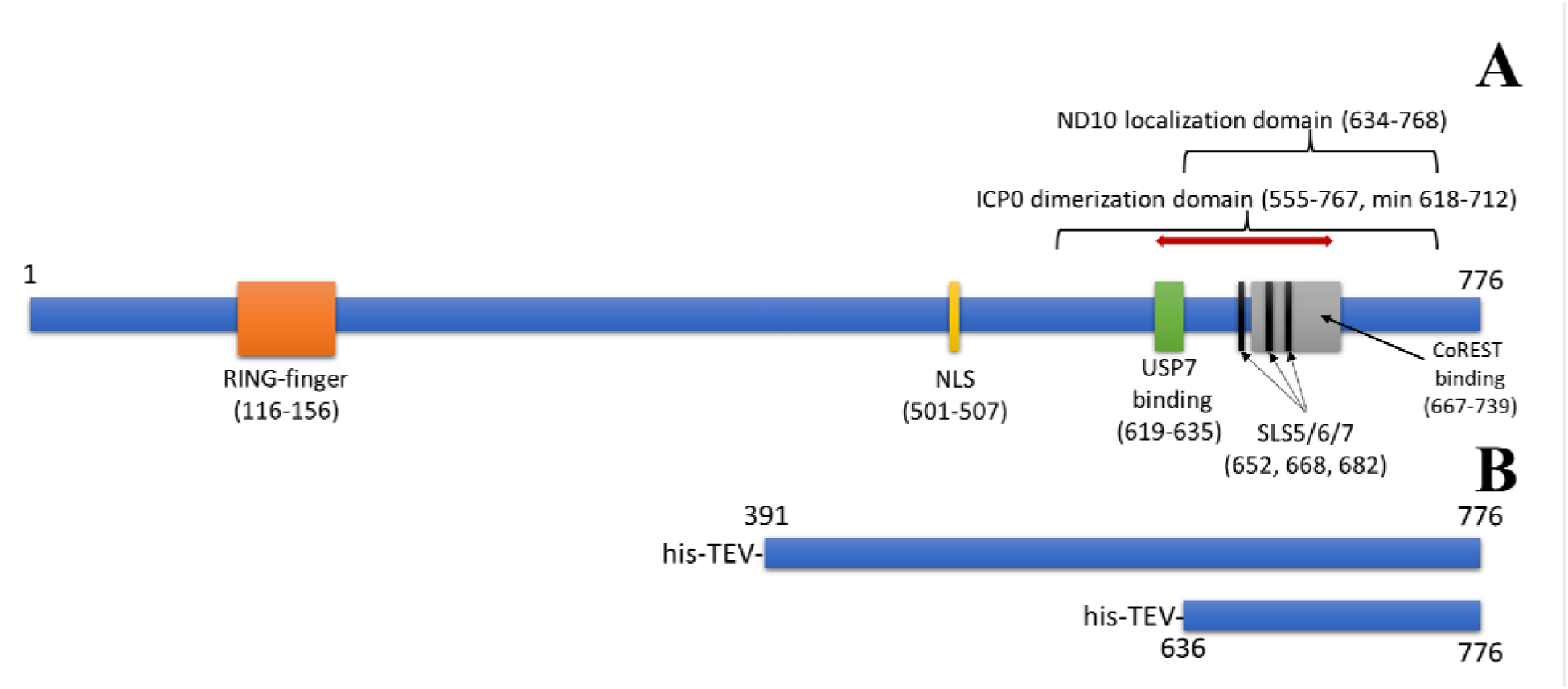
ICP0 Primary Structure from HSV-1 strain KOS. **A)** Important domains indicated. RING-finger domain, residues 116-156. Nuclear Localization Sequence (NLS), residues 501-507. USP7 binding domain, residues 619-635. SUMO-interacting motif (SIM)-like sequences (SLS) 5, 6, and 7: residues 652-655 (SLS5), 668-671 (SLS6), and 682-685 (SLS7). CoREST binding domain, residues 667-739. ND10 localization domain, residues 634-768. ICP0 dimerization domain, residues 555-676, with a minimal dimerization domain from 618-712 (red line). **B)** His-tagged constructs used for structure analyses.

As ICP0 is known to transactivate all classes of HSV genes^7^, its C-terminus is involved in this function. However, it is unclear mechanistically how ICP0’s C-terminus, which contains its dimerization domain, stimulates viral gene expression. Notably, structural studies examining HSV-1 ICP0 have been limited. One study examined the specific motif (∼10-20 amino acids) in ICP0 that interacts with the cellular protein USP7^16^. Another study examined SUMO1 bound to a phosphorylated SLS4, a structure of about 20 amino acids in length^24^. Aside from these studies for ICP0 in HSV-1, the only other large structure that has been described related to ICP0 is its RING-finger domain ortholog from equine herpesvirus^25^. The goal of this present study is to provide the first large structure (>100 amino acids) of HSV-1 ICP0’s C-terminal dimer region using crystallography and X-ray diffraction. We show that the dimer contains two unique small β-barrel structures, and the dimer sequence is highly conserved in non-human primates. Additionally, these structural studies further support the hypothesis that ICP0 is capable of forming tetramers or multimers. Since small β-barrels are highly diverse in their functions, implications of how this unique structure possibly affects functions of ICP0 will be discussed.

## MATERIALS AND METHODS

### Sequence analyses

The sequence of HSV-1 (KOS strain) was obtained from Uniprot and subjected to a BlastP search via the NCBI server^26^. Multiple sequence alignments were generated using COBALT^27^. Additionally, COBALT was also used to align the sequences from various laboratory strains and clinical isolates of HSV-1^28^.

### Expression and Purification of Protein Constructs

Figure 1B shows the his-tagged sequences used for crystallography analyses relative to the primary sequence. The plasmids pTBSG-ICP0 636-776 and pTBSG ICP0 391-776 (based on DNA sequences of HSV-1 strain KOS) were each transformed into BL21 (DE3) pRARE cells and grown overnight at 37°C on LB agar plates with 100 µg/mL ampicillin and 34 µg/mL chloramphenicol. A single colony was added to a flask containing 40 mL LB media, 100 µg/mL ampicillin, and 34 µg/mL chloramphenicol, then incubated overnight at 37°C with shaking. Overnight culture was divided equally and added to two shaker flasks containing 2 L LB media, 100 µg/mL ampicillin, and 34 µg/mL chloramphenicol, then shaken at 200 rpm at 37°C until OD600 of 0.7. A final concentration of 0.4 mM IPTG was added and induced for 2 hours at 37°C with shaking. Cells were pelleted at 3500 rcf for 10 minutes, and the supernatant was discarded. The cell pellet was resuspended in 50 mM Tris (pH 8.0) and 500 mM NaCl and frozen at −80°C.

Frozen lysate was thawed and sonicated until it was no longer viscous, then spun at 45,450 rcf for 30 minutes at 4°C. The supernatant was decanted to isolate the soluble fraction, and the insoluble pellet was resuspended in water for SDS-PAGE gel analysis. The soluble fraction was purified by Ni-NTA affinity chromatography using AKTA Purifier and a 5 mL HisTrap HP column prepacked with Ni Sepharose High Performance beads. The whole cell lysate, soluble fraction, insoluble fraction, flowthrough, and three purified fractions were analyzed on a 12% SDS-PAGE gel and stained with InstantBlue Protein Stain. The purified fractions were pooled, and 6 mL was reserved for size exclusion chromatography (SEC). The remaining pooled protein was concentrated to 10 mL by centrifugal membrane filtration (MWCO 3 kDa). Glycerol was added to 50% and saved at −20°C.

The NiNTA-purified larger construct (his-ICP0 391-776) was also subjected to TEV protease digestion with his-TEV (0.6 mg) at 4°C overnight in a dialysis tubing, dialyzed against 1 L 50 mM Tris (pH 8.0), 500 mM NaCl.

SEC was performed using AKTA Purifier and Superdex 200 10/300 GL column, with an injection volume of 2 mL. Buffer 50 mM Tris (pH 8.0), 250 mM NaCl. Protein factions were analyzed on a 12% SDS-PAGE gel stained with InstantBlue Protein Stain. Fractions from each run were pooled for a final volume of 12 mL and concentrated to roughly 0.6 mL by centrifugal membrane filtration (MWCO 5kDa). Final concentration was determined by Nanodrop at A280nm.

### Crystallization and Data Collection

Purified ICP0 containing the purification tag was concentrated to 10.4 mg/mL in 250 mM NaCl and 50 mM Tris pH 8.0. All crystallization experiments were set up using an NT8 drop setting robot (Formulatrix Inc.) and UVXPO MRC (Molecular Dimensions) sitting drop vapor diffusion plates at 18°C. 100 nL of protein, and crystallization solution (100 nL) was dispensed and equilibrated against 50 µL of the latter. Crystals that displayed a prismatic morphology (**Supplemental** Figure 1) were observed after approximately 2 weeks and grew to their maximum size (∼100 µm x 50 µm) over two months from the PACT screen (Molecular Dimensions) condition G3 (20% (w/v) PEG 3350, 100 mM Bis-Tris Propane pH 7.5, 200 mM NaI). A cryoprotectant solution composed of 80% crystallization solution and 20% (v/v) PEG 200 was dispensed (2 µL) onto the drop, and crystals were harvested immediately and stored in liquid nitrogen. X-ray diffraction data were collected at the Advanced Photon Source beamline 17-ID using a Dectris Pilatus 6M pixel array detector. Two data sets were collected using the same crystal at wavelengths of 1.0000 Å (HE) and 1.5895 Å (LE). The LE data set was collected to enhance the anomalous signal for phasing in the event that iodide ions from the crystallization solution were bound to the protein, and the HE data were used for final refinement.

The first, larger construct spanning residues 391-776 failed to form usable crystals. From this, a proteolytic fragment was able to crystalize and the second construct was created to match this proteolytic fragment. This second construct spanning residues A636-Q776 (His-ICP0 in **Supplemental Table 1**) that contained a 6X-His tag and TEV protease site was concentrated to 16 mg/mL in the same buffer as above. Crystals for data collection were obtained from the PACT screen (Molecular Dimensions) condition H3 (20% (w/v) PEG 3350, 100 mM Bis-Tris Propane pH 8.5, 200 mM NaI) and were cryoprotected as noted above.

### Structure Solution and Refinement

Intensities were integrated using XDS^29,30^ via Autoproc^31^ and the Laue class analysis and data scaling were performed with Aimless^32^. Structure solution was conducted by SAD phasing with Crank2^33^ using the Shelx^34^, Refmac^35^, Solomon^36^, Parrot^37^, and Buccaneer^38^ pipeline via the CCP4^39^ interface using the LE data set. Two highly occupied iodide ions were located (1.0 and 0.75 relative occupancies) and phasing/density modification resulted in a mean figure of merit of 0.47. Subsequent model building with density modification and phased refinement yielded *R*/*R*_free_ = 0.41/0.50 for the refined model. This model was then used for molecular replacement with Phaser^40^ against the higher resolution HE data set, and the top solution was obtained in the space group *P*4_1_2_1_2 (RFZ=11.2 TFZ=26.3 LLG=562). The model was further improved by automated model building within the Phenix^41^ suite. Additional refinement and manual model building were conducted with Phenix and Coot^42^, respectively. The crystals obtained from the A636-Q776 (His-ICP0) construct were isomorphous with the original crystals obtained from PACT G3, and solution was obtained by molecular replacement using the model described above. Disordered side chains were truncated to the point for which electron density could be observed. Structure validation was conducted with Molprobity^43^, and figures were prepared using the CCP4MG package^44^. Crystallographic data are provided in **Supplemental Table 1**.

### Molecular modeling of ICP0

Monomeric models of ICP0 were generated using Alphafold 2.2 using the sequence of HSV-1 (KOS Strain) ICP0^45^. Of the five models generated, the model with the highest pLDDT (44.8) was selected for analysis. Tetramers of the ICP0 C-terminal domain (637-776) were generated using Alphafold-multimer 2.2^45,46^. Of the 25 models generated, the top model based on ipTM+pTM (0.457) was selected for analysis.

### Molecular modeling of SUMO1-ICP0 complex

Models of SUMO1 bound to the SLS5 and SLS7 of ICP0 were generated using a template-based approach, as SUMO1-bound structures in the PDB share a common binding to SLS domains. Because SUMO1 has been observed binding both parallel and antiparallel to its various targets, two different templates were used: ICP0 phosphorylated SLS4 (6JXV^24^) as the parallel template and RanBP2 (2LAS^47^) as the antiparallel template. Monomers from the experimentally determined ICP0 tetramer were used, however, because of potential flexibility of monomeric ICP0. Only the secondary structure elements that were associated with each SLS were modeled. For SLS5, residues 640-656 formed part of a five-stranded, twisted β-sheet along with residues 730-755. For SLS7, residues 666-711 formed a β-hairpin connected to two helixes.

The four-residue SLS segment from each of the two SLS-containing substructures was aligned to the four-residue SUMO-interacting residues of each of the two templates. SLS5 was only capable of binding SUMO1 in an antiparallel manner, while SLS7 was only capable of binding it in a parallel manner. Each of the resulting models were subjected to one thousand independent simulations using Rosetta Relax^48^ and the top scoring complexes were chosen as the final models. Comparison of the energy of the complex to the combined energy of unbound SUMO1 and the SLS-containing substructure (-320.223 vs - 287.773 Rosetta Energy Units for SLS5, -338.013 vs 291.849 REUs for SLS7) suggested that complex formation is, indeed, energetically favorable.

## RESULTS

### Sequence Analyses

The sequence of full length HSV-1 ICP0 was subjected to a BLAST search against the RefSeq database. It became immediately apparent that the CTD was not conserved across all ICP0 orthologs from alphaherpesviruses; it was only present in alphaherpesviruses of primates, with the exception of varicella-zoster virus (VZV) or simian varicella-zoster virus, which lacks the CTD. The CTD-containing viruses have varying degrees of conservation. For example, when looking at the multiple sequence alignment of the whole CTD, viruses targeting new world monkeys (Saimiriine herpesvirus-1 and Ateline herpesvirus-1) have noticeably less conservation (Figure 2A). Many of the residues between L644 and D686 make up the dimeric interface, which correlates to the region with the most sequence conservation (Figure 2B). In fact, of the 27 residues within 3 Å of binding to the other chain in the dimer, 12 are fully conserved (P645, S651, T662, G665, D666, C667, P669, D672, I678, Y681, V683, and V685). An additional 12 residues are conserved in herpesviruses targeting old world monkeys, apes, and humans (S650, V652, A654, Y658, N660, K661, T664, L671, G676, G679, V716, and H760). Residues S647, M673, and E674 are the least conserved.

**Figure 2:**
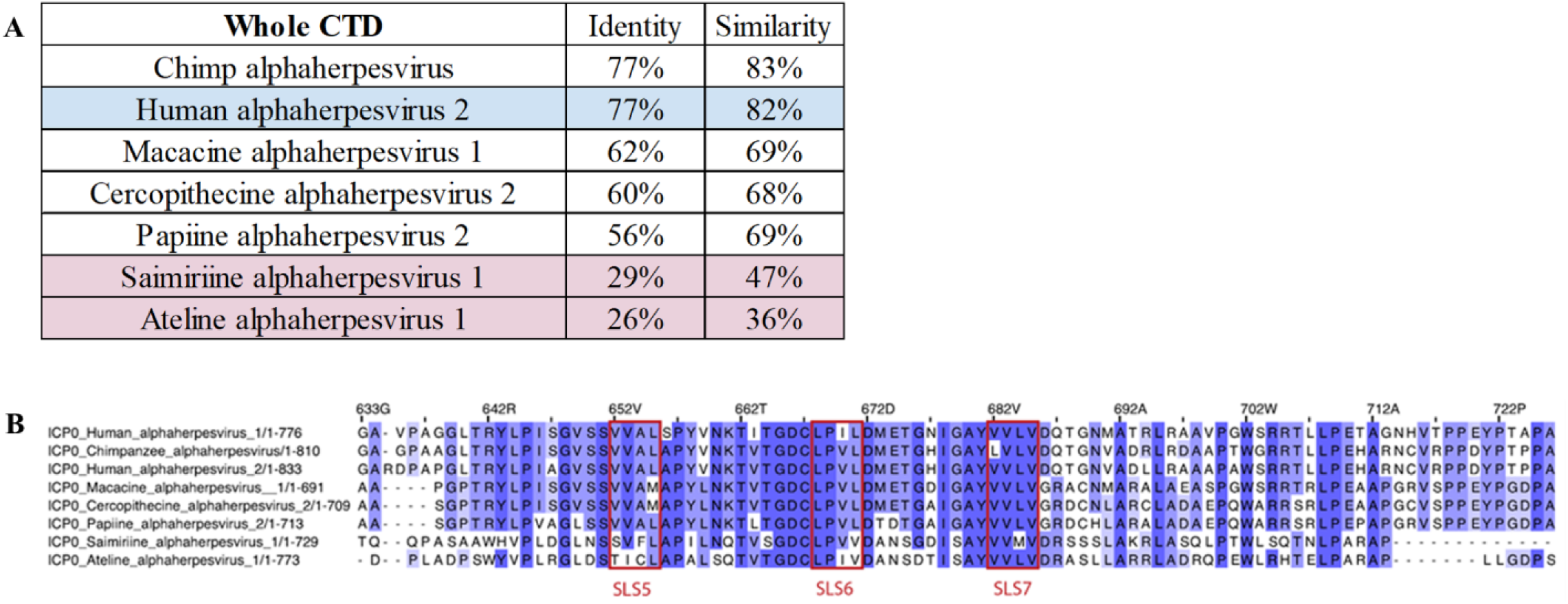
Conservation of residues in the ICP0 CTD. **A)** Identity and similarity of the entire CTD of ICP0 in various primate alphaherpesviruses compared to HSV-1. Blue indicates HSV-2, and red indicates alphaherpesviruses from new world monkeys. **B)** Alignment of ICP0 CTD dimer interface residues. The dimer interface region of ICP0 in unique herpesvirus species containing the CTD. SLS5, SLS6, and SLS7 are marked. The numbering is relative to HSV-1 ICP0.

While the dimeric interface was composed of 27 residues within 3Å, the tetramer (dimer of dimers) interface is much smaller, with only four residues per chain within 3Å of the other dimer. Of those four residues, only S650 and T735 are fully conserved in herpesviruses targeting old world monkeys, apes, and humans. An additional six residues are between 3Å and 3.5Å, with F742 and G745 fully conserved, and W733 and L740 conserved in herpesviruses targeting old world monkeys, apes, and humans. Indeed, there are seven contacting residues between W733 and G745, and that region has more conservation (**Supplemental** Figure 2), though little conservation with herpesviruses targeting new world monkeys. A diverse collection of 16 HSV-1 ICP0 clinical isolates and strains were assembled, and the portion of the CTD that was solved was almost completely conserved; however, one outlier was T717, a solvent exposed residue that is mutated to a methionine in some strains/isolates (**Supplemental** Figure 3).

### Structure Analyses

The his-tagged ICP0 construct spanning A391-Q776 is 41,108 Da and one would expect a monomer to occupy the asymmetric unit based on the Matthews coefficient^49^ (Vm=2.1 / 41% solvent). The electron density maps following phasing were of high enough quality to clearly discern features that were consistent with a polypeptide. However, attempts to model the sequence spanning A391-Q776 to the maps failed. It became clear that the electron density was not consistent with this construct, and 2-fold non-crystallographic symmetry (NCS) appeared to be present in the maps. It was ultimately determined that a smaller fragment of ICP0 had crystallized, and a model composed of an NCS dimer spanning L640-S763 in each subunit was constructed. Therefore, a new recombinant construct was produced that spanned A636-Q776 and contained a 6X-His tag and TEV protease site. These crystals were isomorphous with those observed from that fragmented construct and were used to obtain the structure described here. Residues P722-W729 (subunit A) and E720-N730 (subunit B) were disordered and could not be modeled.

Individual subunits are composed of nine β-strands and two α-helices producing the following arrangement: β1-β2-β3-β4-α1-α2-β5-β6-β7-β8-β9. The spatial arrangement of the β-strands can be envisioned as three groups consisting of an anti-parallel β-barrel-like motif spanning β1-β2-β6-β7-β8 (βG1), a long pair of anti-parallel strands at β3-β4 (βG2) and a short pair of anti-parallel strands at β5-β9 (βG3). Helices α1-α2 bridge the β-strands between βG2 and βG3. The structure of subunit A is depicted in Figure 3A as a ribbons diagram, and a topology diagram showing this arrangement is provided in Figure 3C. Annotations of these structural elements relative to sequence are shown in Figure 3B. Hydrogen bonds between β-strand groups are listed in **Supplemental Table 2**.

**Figure 3:**
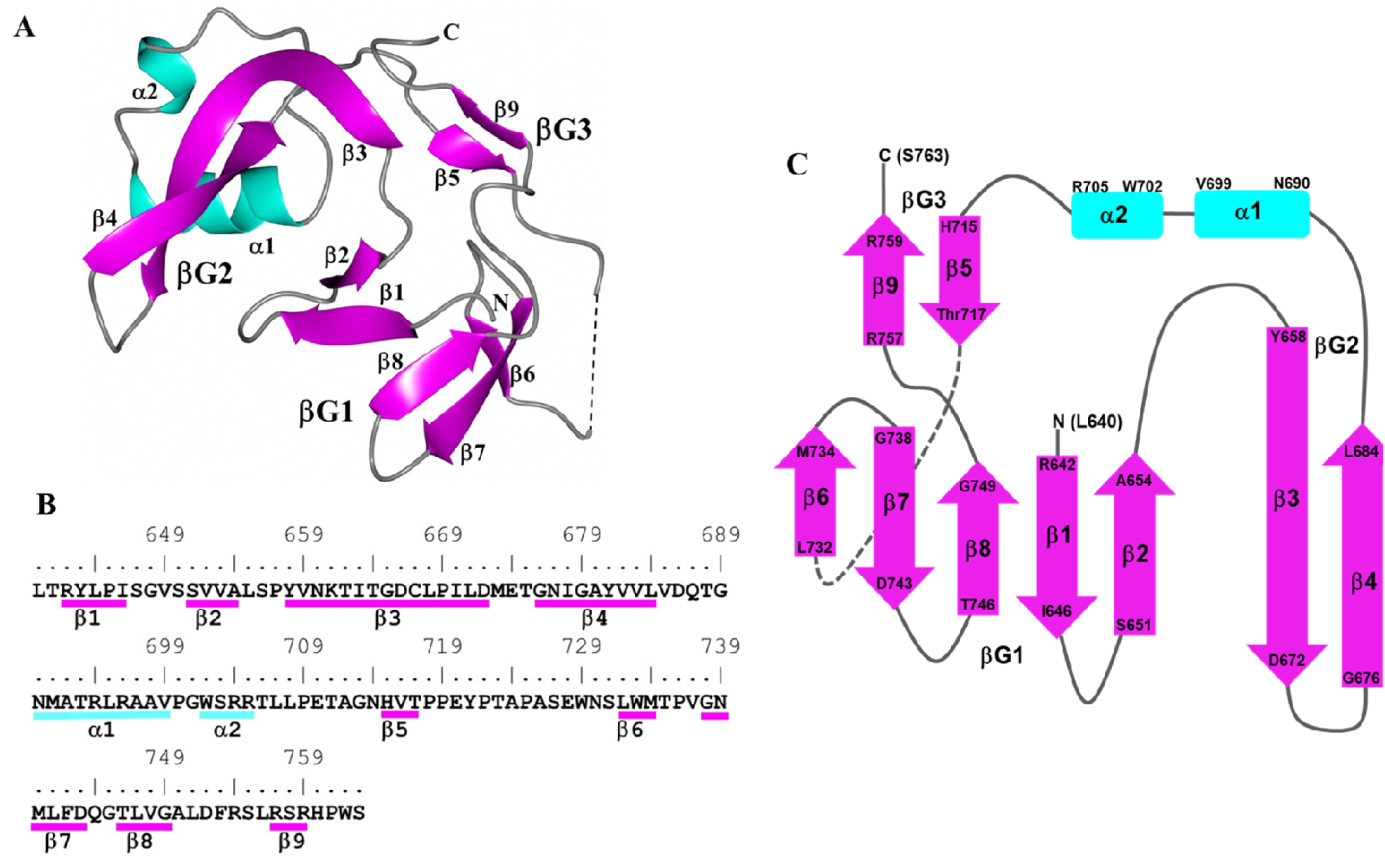
Subunit A of ICP0. **A)** Ribbon diagram of secondary structures with helices (cyan) and β-strands (magenta) indicated. The dashed line indicates a break in the chain between Y721 and N730 due to disorder. N and C correspond to L640 and S763 at the terminal ends. **B)** Secondary structure elements are annotated relative to the ICP0 sequence. **C)** Topology diagram of ICP0 showing the arrangement of the three β-strand regions (βG1-3) and two α-helices.

Further analysis of subunit A reveals a cavity that is formed by βG2, βG3, a portion of the β4 strand and α1 (Figure 4A and C). Formation of the non-crystallographic dimer of ICP0 results in an interaction between β3 and its counterpart in subunit B (β3’), which creates an intertwined β-strand pair (Figure 4B). As such, strands β3’ and β4’ in subunit B insert into the aforementioned cavity as depicted in Figure 4D. This results in new interactions between strands β3-β3’ and β4-β4’ located at the center of the dimer (Figure 4E). In addition, β3’ and β4’ of subunit B are positioned in the cavity in subunit A between β2 and β5, generating a nine-strand β-barrel-like structure (Figure 4F), in which β6 and β9 overlap parallel to one another like a single strand. Hydrogen bond interactions between subunits A and B are listed in **Table 1**. A more detailed structure indicating some of the hydrogen bonds between the subunits can be visualized in **Supplemental** Figure 4. As noted previously, two iodide ions were modeled. These ions are located near helix α1 as shown in **Supplemental** Figure 5 along with the anomalous difference in electron density.

**Figure 4:**
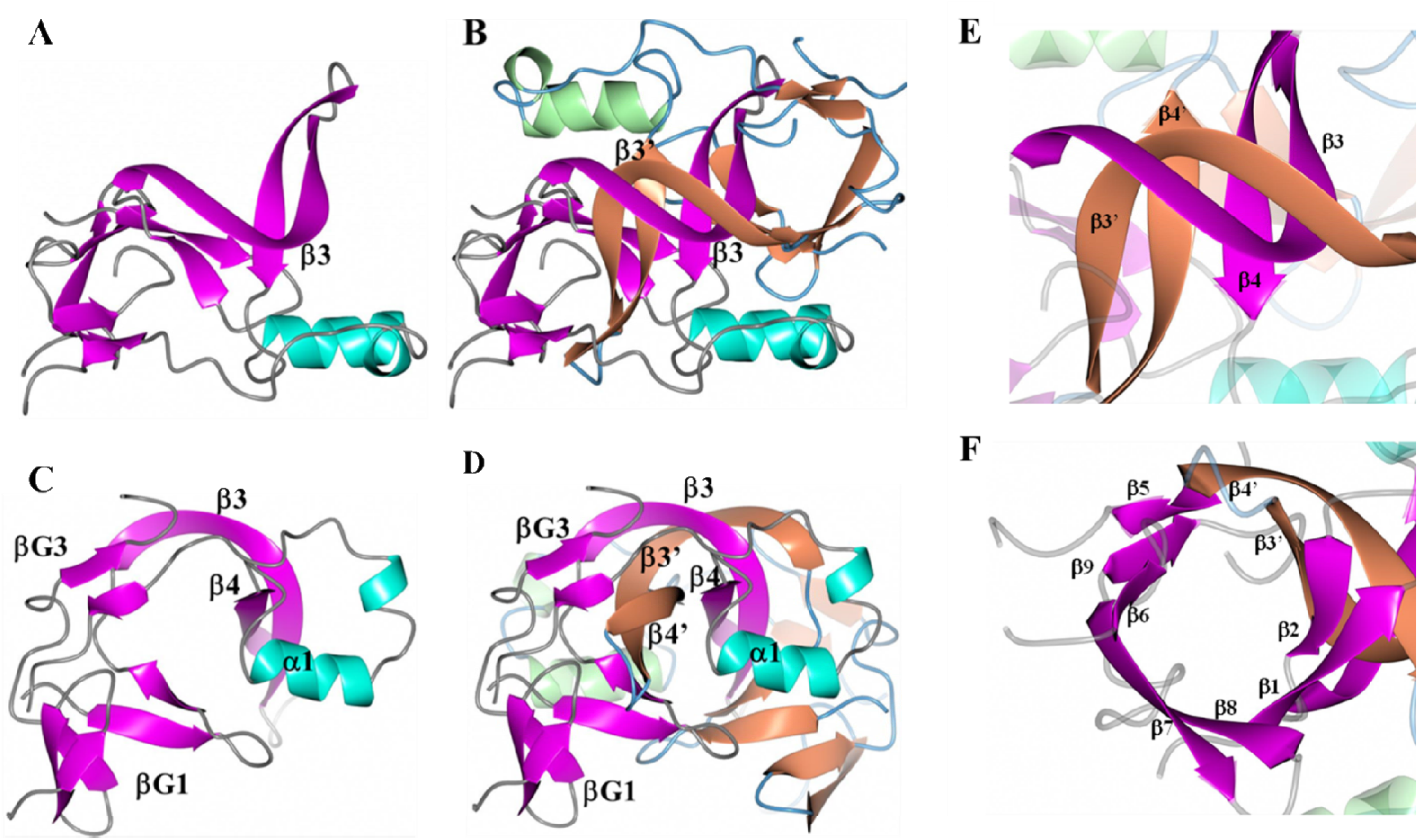
ICP0 dimer arrangement. **A)** Subunit A showing helices (cyan), β-strands (magenta), and loops (gray). **B)** Subunit B added with helices (green), β-strands (tan), and loops (blue). The β3 strand intertwines with its counterpart, β3’. View is along the non-crystallographic 2-fold axis. Panels **C** and **D** are views perpendicular to the non-crystallographic 2-fold axis. **E)** Anti-parallel β-strand interactions are formed between β3-β3’ and β4-β4’. **F)** β3’ and β4’ of subunit B are inserted between β2 and β5 of subunit A to form a nine-strand β-barrel-like structure.

**Table 1:**
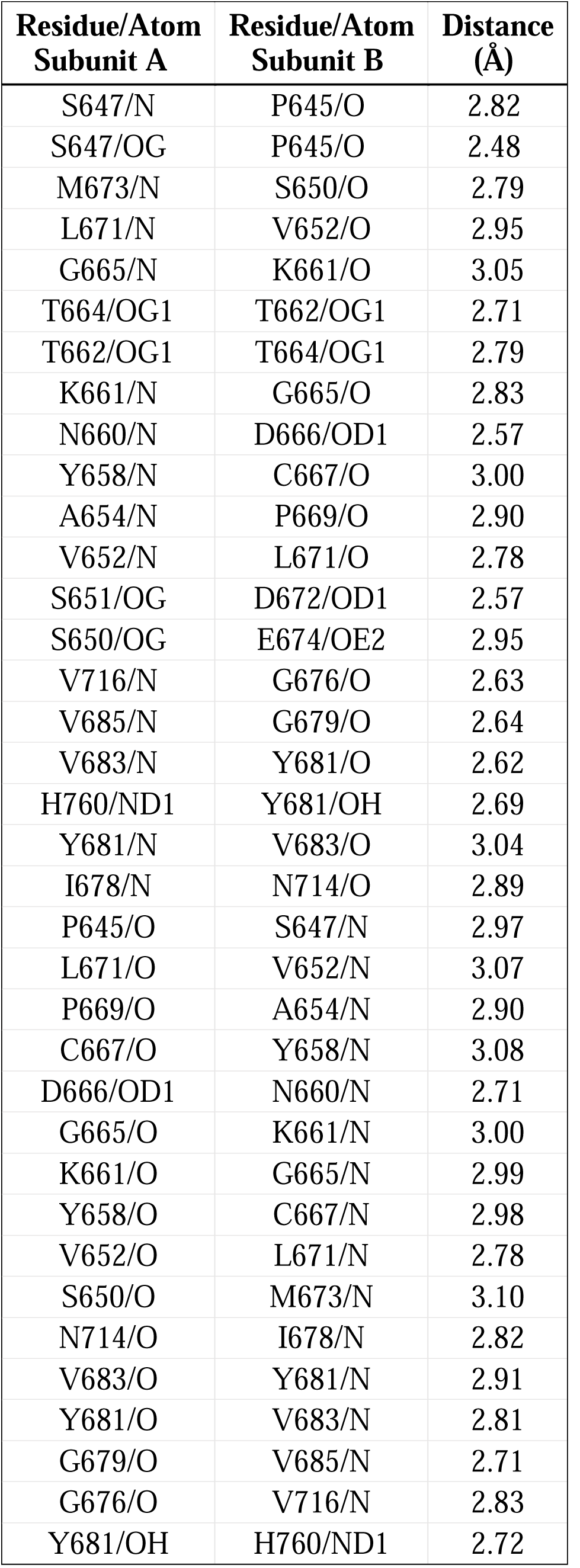
Hydrogen bond interactions between subunits A and B of ICP0.

| **Residue/Atom**<br>**Subunit A** | **Residue/Atom**<br>**Subunit B** | **Distance**<br>**(Å)** |
| --- | --- | --- |
| S647/N | P645/O | 2.82 |
| S647/OG | P645/O | 2.48 |
| M673/N | S650/O | 2.79 |
| L671/N | V652/O | 2.95 |
| G665/N | K661/O | 3.05 |
| T664/OG1 | T662/OG1 | 2.71 |
| T662/OG1 | T664/OG1 | 2.79 |
| K661/N | G665/O | 2.83 |
| N660/N | D666/OD1 | 2.57 |
| Y658/N | C667/O | 3.00 |
| A654/N | P669/O | 2.90 |
| V652/N | L671/O | 2.78 |
| S651/OG | D672/OD1 | 2.57 |
| S650/OG | E674/OE2 | 2.95 |
| V716/N | G676/O | 2.63 |
| V685/N | G679/O | 2.64 |
| V683/N | Y681/O | 2.62 |
| H760/ND1 | Y681/OH | 2.69 |
| Y681/N | V683/O | 3.04 |
| I678/N | N714/O | 2.89 |
| P645/O | S647/N | 2.97 |
| L671/O | V652/N | 3.07 |
| P669/O | A654/N | 2.90 |
| C667/O | Y658/N | 3.08 |
| D666/OD1 | N660/N | 2.71 |
| G665/O | K661/N | 3.00 |
| K661/O | G665/N | 2.99 |
| Y658/O | C667/N | 2.98 |
| V652/O | L671/N | 2.78 |
| S650/O | M673/N | 3.10 |
| N714/O | I678/N | 2.82 |
| V683/O | Y681/N | 2.91 |
| Y681/O | V683/N | 2.81 |
| G679/O | V685/N | 2.71 |
| G676/O | V716/N | 2.83 |
| Y681/OH | H760/ND1 | 2.72 |

Analysis of the crystal packing indicated that a tetrameric arrangement, with 222 point symmetry, was present by applying the 2-fold crystallographic symmetry operator (Y, X, Z) on the asymmetric unit. The likelihood of higher order assemblies of ICP0 was examined with Pisa^50^, which also indicated that a tetramer was the most likely arrangement for ICP0, depicted in Figure 5A. Notably, the tetrameric assembly contains a hole in the middle that is approximately 11 Å high x 9 Å wide and 15 Å deep. The main interactions between the dimers to form the tetramer occur between β6 and β7 of the symmetry-related dimer, which adopt a parallel orientation (Figure 5B). Hydrogen bond interactions between dimers in the tetramer are listed in **Supplemental Table 3**. Interestingly, the formation of the tetramer extends the β-barrel-like motif. This can be envisioned as the stacking of β-barrels from each of the dimer units, by the pairing of β6 and β7, which creates an extended tunnel as shown in Figure 5C. The electrostatic surface of the tetramer contains patches of negative and positive charge on the “top” of the assembly as depicted in Figure 5D. It should be noted that the monomer, dimer, and tetramer forms of ICP0 were subjected to searches using the DALI and PDBeFold servers. The top match was obtained with PDB 5A1U (CopI Coat Protein) using the dimer of ICP0 as the search template. However, only 39 residues could be aligned and the RMSD deviation was 3.72Å, indicating a poor fit. As such, ICP0 appears to adopt a novel structure/fold.

**Figure 5:**
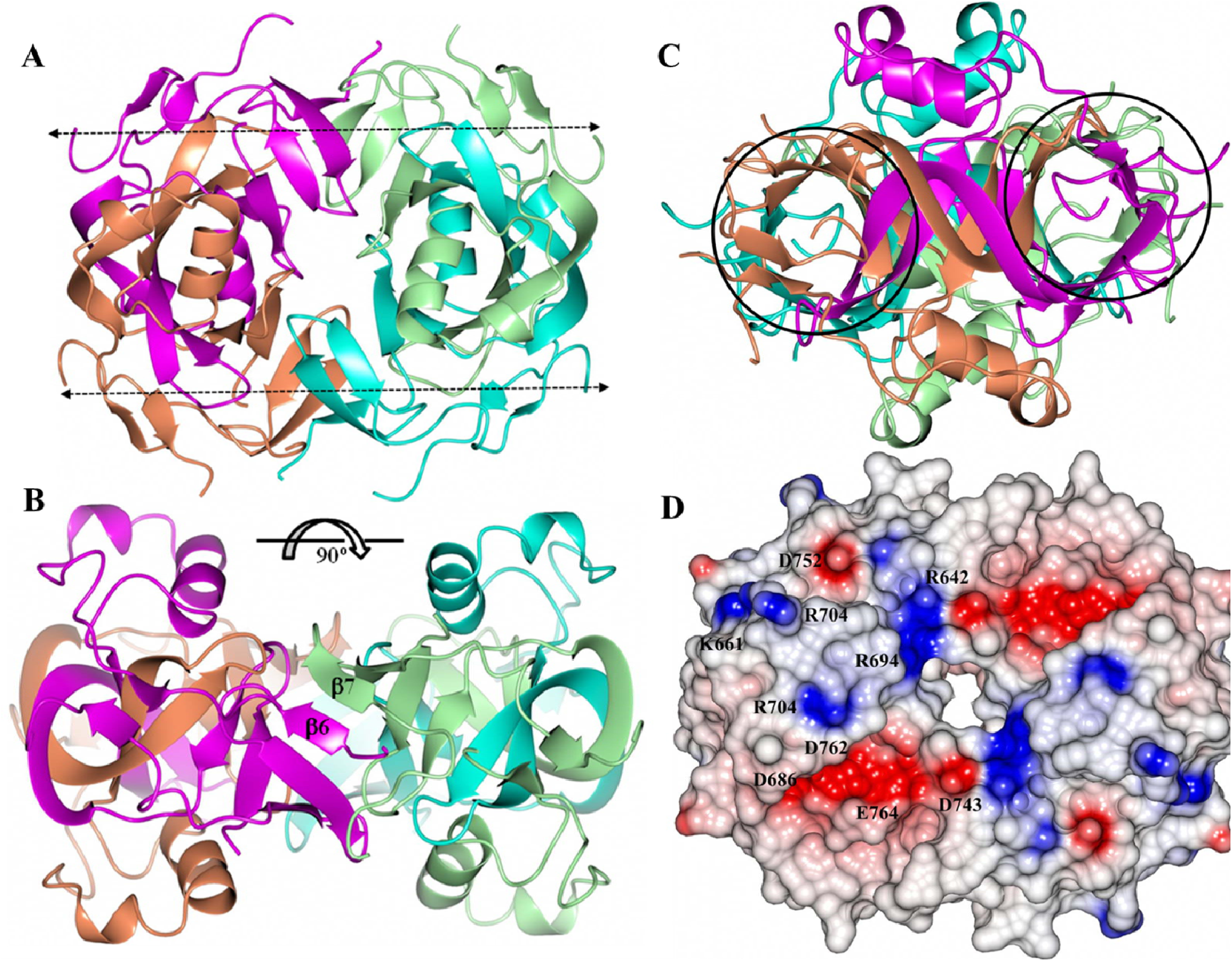
ICP0 tetramer assembly with subunits colored A (magenta), B (tan), C (cyan) and D (green). **A)** View along the crystallographic 2-fold axis. The dashed arrows indicate the direction of the β-barrel-like motifs as viewed in Figure 4B. **B)** View perpendicular to the crystallographic 2-fold axis. The β6 and β7 strands which form hydrogen bond interactions are indicated. **C)** The regions in circles are the extended β-barrel-like motifs viewed along the barrel axis. **D)** Electrostatic surface of the ICP0 tetramer with the residues contributing to the positive/negative patches indicated. The scale is from -0.5 V (red) to 0.5 V (blue).

One feature that stood out in the crystal structure was the nine-strand β-barrel-like structure within the dimer, which stack in the tetrameric form. They were reminiscent of small β-barrels, which were reviewed by Youkharibache et al.^51^, although they had a different topology. The ICP0 CTD β-barrels have a roughly conical shape (Figure 6A), with one end having backbone-backbone distances between 12-16 Å, 10-12 Å on the other, and are roughly 12-13 Å tall. Several small β-barrels were selected from the survey by Youkaribache et al. and are also roughly conical^51^. The Hfq β-barrel, 1KQ2^52^, (Figure 6B) is slightly wider and shorter, with one end 14-18 Å wide and the other 13-16 Å, and 9-11 Å tall. The shiga-like toxin B β-barrel, 1C4Q, (Figure 6C) is slightly wider and about the same height, with one end 15-17 Å wide and the other 10-13 Å, and has a height of 10-13 Å. The *Drosophila* HP1 chromodomain and histone H3 tail complex to form a β-barrel, 1KNA^53^ (Figure 6D), which is composed of two chains and is less conical and shorter, with one end 12-14 Å wide, the other 12-13 Å, and a height of 9-11 Å. Thus, small β-barrels may be composed of one or two chains, tend to be wider at one end, with the three mentioned between 12-18 Å wide at one end and 10-16 Å at the other, and heights ranging around 9-13 Å tall. The ICP0 small β-barrel has similar properties to these other small β-barrels and appears to represent a previously unknown small β-barrel topology.

**Figure 6:**
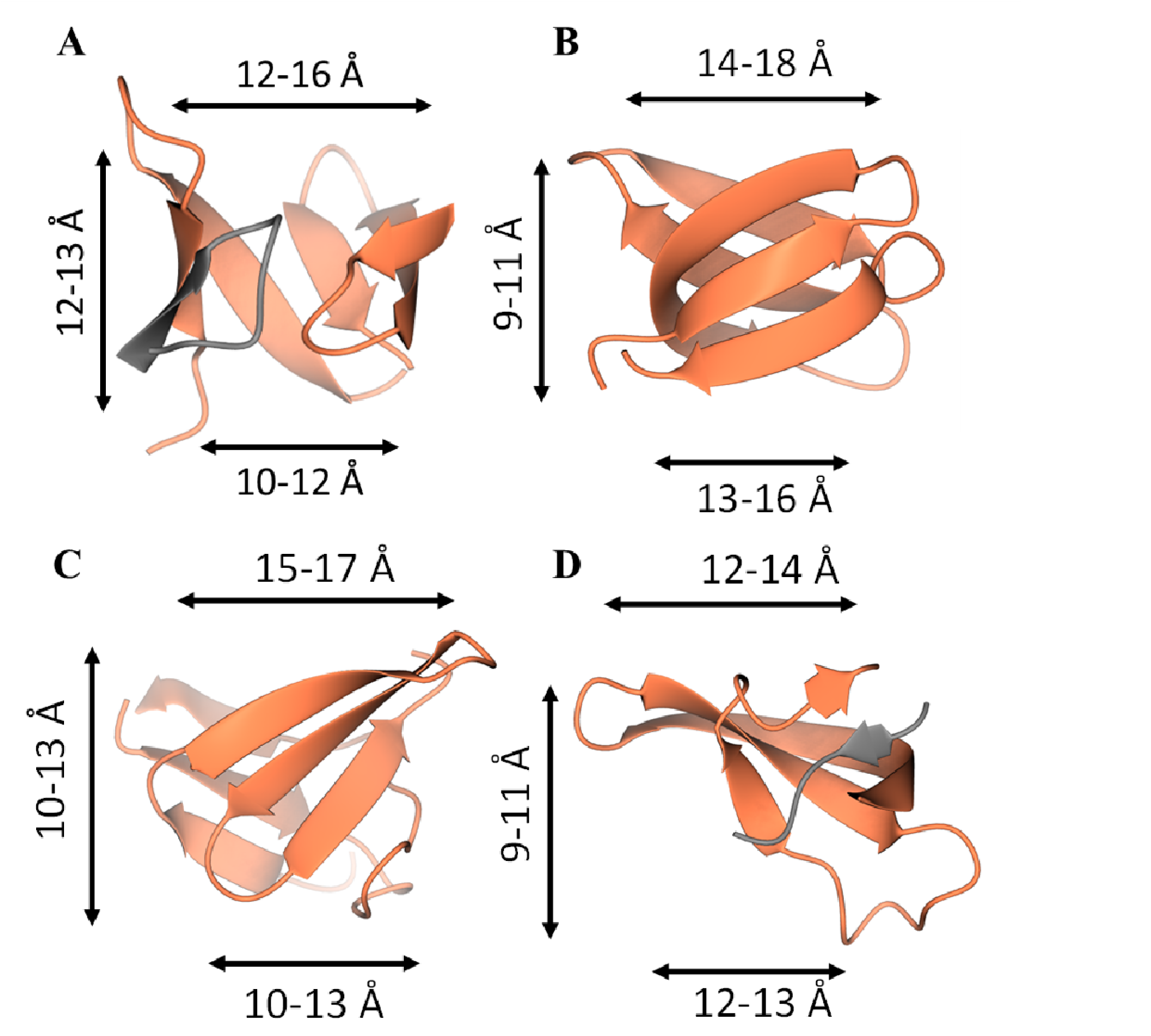
Four small β-barrels. **A)** The ICP0 CTD β-barrel. The two chains comprising the β-barrel are isolated and colored by chain, with the narrower end of the β-barrel on the bottom. **B)** The Hfq β-barrel, with the narrower end of the β-barrel on bottom. **C)** The shiga-like toxin B β-barrel with the narrower end of the beta barrel on bottom. **D)** The **Drosophila** HP1 chromodomain and histone H3 tail complex. The two chains comprising the β-barrel are isolated and colored by chain, with the narrower end of the β-barrel on bottom. Ranges of Cα-Cα distances are marked.

### Computational Predictions of the CTD

Because the ICP0 CTD adopts a novel conformation, we sought to assess how well Alphafold^45^, the current state of the art tool for protein structure prediction, predicted this novel fold. First, the full-length structure was predicted as a monomer. Most of the structure was disordered, aside from the CTD and a portion of the N-terminal domain (NTD).

The CTD residues 639-776 were modeled using Alphafold as both a monomer and the tetramer. The model of the CTD monomer had an average predicted local distance difference test (pLDDT) of 49, suggesting a model of very low confidence. At the per-residue level, much of the predicted structure has very low confidence, with pLDDT values under 50, yet D666 to A712 is predicted with somewhat higher confidence to have a folded structure, with 15 residues having pLDDT values exceeding 70 (**Supplemental** Figure 6). This region included two strands and a helix. Superposition of this region onto the crystal structure shows that the prediction of residues 666-712 as a monomer was quite accurate, with a 0.96 Å Cα RMSD (**Supplemental** Figure 7).

Alphafold-multimer^46^ was also able to generate a model of the CTD that resembled aspects of the crystallographic structure, with a Cα RMSD of 2.80 Å. As seen in **Supplemental** Figure 8, some portions modeled virtually identical to the crystal structure, while other regions look nothing like the crystal structure. For example, residues G748-713, which form the core of the dimer, were reasonably well predicted (see the left and right side of each tetramer in the top panel (**Supplemental** Figure 8A and B) and the center of the tetramer in **Supplemental** Figure 8). The rest of the molecule, including the tetramer interface and half of the strands that compose the stacked β-barrels (**Supplemental** Figure 8C and D), did not model well compared to the crystal structure. This matches the pLDDT of the predictions, with P657-G713 having pLDDT values greater than 0.5, and the central core of the dimer interface residues having pLDDT values greater than 0.7 (**Supplemental** Figure 9). Of particular note, Alphafold essentially makes structural predictions based on predicted contact networks inferred from sequence covariance of homologous proteins across species. The tetramer interface, where the prediction does not match most of the crystal structure, is the region that differs between new world monkeys and the rest of the primates. This has implications for both the evolution of ICP0 tetramerization (associated with the sequencing and annotation of additional viral genomes) and increasing the accuracy of predicting novel structural domains in viral proteins.

### SLS5/6/7 and SUMO1

The CTD of ICP0 contains three of the seven SUMO-binding domains in ICP0: SLS5, SLS6, and SLS7. Visual inspection of the solved ICP0 CTD shows that SLS5-7 are located at the dimeric interface (Figure 7A). When looking at only one chain, the SLS appear to be solvent exposed (Figure 7B), yielding the question of whether or not they can interact with SUMO1 when present as a monomer, particularly SLS5 and SLS7. SUMO1 tends to bind proteins in such a way that edge of the SUMO1 β-strand is extended by its binding partner. The binding partner can interact either parallel or anti-parallel to the SUMO1 β-strand, depending on the protein. Because there are flexible, extended regions between the β-strands containing SLS5 and SLS7, the folded regions of the monomers were treated separately and docked to SUMO1 in both the parallel (6JXV^24^), and anti-parallel (2LAS^47^) conformations. SLS5 and SLS7 were aligned to other SIMs in complex with SUMO1. Feasible models were then refined using Relax in Rosetta^48^.

**Figure 7:**
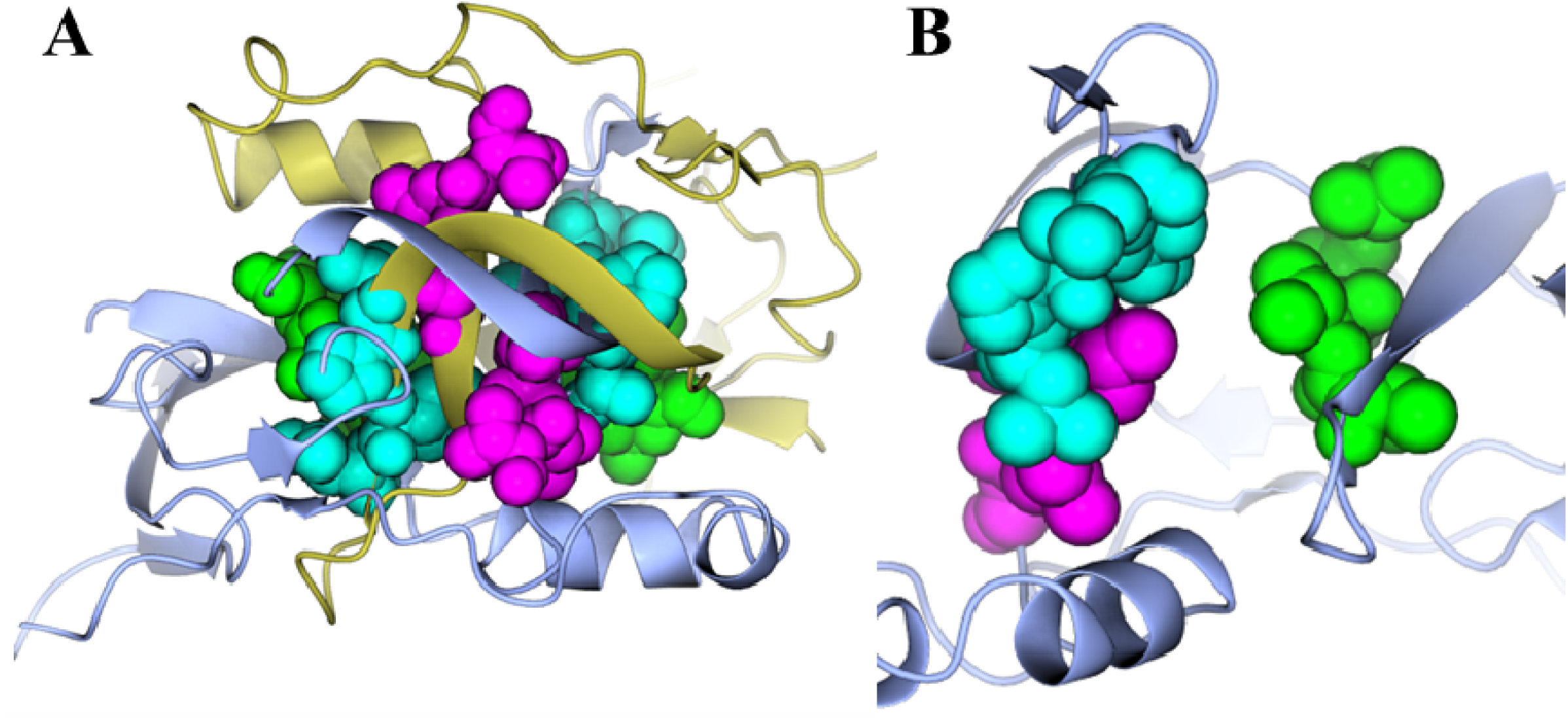
SLS5, SLS6, and SLS7 are located at the dimeric interface. **A)** The two chains comprising the dimer (each chain is a different color) are shown as ribbons. SLS5 (green), SLS6 (cyan), and SLS7 (magenta) are shown as spheres. **B)** A single chain is shown as ribbons, while SLS5 (green), SLS6 (cyan), and SLS7 (magenta) are shown as spheres.

A model of SLS5 bound to SUMO1 was created, which was only capable of binding to SUMO1 in an anti-parallel manner (**Supplemental** Figure 10). Energetically, the top scoring complex was more favorable than the sum of each monomer, with a score of -320.223 Rosetta energy units (REU), compared to the less favorable scores of -214.334 and -77.772 for SUMO1 and ICP0, respectively. Additionally, a model of SLS7 bound to SUMO1 was also created, only capable of binding SUMO1 in a parallel manner (**Supplemental** Figure 11). While not definitive, the model-driven hypothesis of interactions between SUMO1 and SLS5 and SLS7 warrants further investigation. Taken together, these models suggest that ICP0 monomers can form dimers or ICP0 monomers can interact with SUMO1 but cannot do both.

## DISCUSSION

In this study, we sought to characterize the structure of the ICP0 C-terminal dimerization domain. Purified ICP0 was crystallized and assessed using X-ray diffraction. Subunits were found to be composed of nine β-strands and two α-helices (β1-β2-β3-β4-α1-α2-β5-β6-β7-β8-β9). The first group of this arrangement (β1-β2-β6-β7-β8), collectively called βG1, forms an anti-parallel β-barrel-like motif. The second group, βG2 (β3-β4), forms a long pair of anti-parallel strands, and the third group, βG3 (β5-β9), forms a short pair of anti-parallel strands. The helices (α1-α2) bridge βG2 and βG3. During dimer formation, β3 and β4 and their counterparts in the opposite subunit, β3’ and β4’, create intertwined pairs, generating a nine-strand β-barrel-like structure. Further analysis indicated the formation of a tetramer that stacks and extends the β-barrels by pairing β6 and β7. The monomer, dimer, and tetramer forms of ICP0 were each compared to other proteins and the fold determined to be novel. Additional analysis with Alphafold revealed that residues 666-712 of the C-terminal domain were correctly predicted when a monomer, with a Cα RMSD of 0.96 Å.

This article highlights the first large structural analysis of the ICP0 protein in HSV-1. Despite the identification of five HSV-1 immediate-early proteins, there are limited models describing their structures. Consequently, this study adds to our knowledge of the HSV-1 IE proteins. Other than USP7^16^ and phosphorylated SLS4 bound to SUMO1^24^, the only other large structure of an ICP0 ortholog that has been described is the structure of the RING-finger domain^25^. ICP27 is another HSV-1 IE protein that plays major roles in post-transcriptional viral gene expression^54^. The C-terminal domain of ICP27 forms a homodimer of α-helical bundles, akin to the dimer structure described here for ICP0, although the β-barrel motif of the ICP0 dimer appears to be unique^54^. Studies of ICP27 homologues suggest that mutations in the dimer domain “tail motifs” lead to loss of the ability to dimerize and perform its transactivation functions^54^. It is possible a similar domain effect may be observed for ICP0 dimerization mutants.

The C-terminal domain described herein does not exist in non-primate herpesviruses, and appears to be conserved in non-human primates, although the alphaherpesviruses that infect new world monkeys have less similarity. The region within ICP0 that contains the highest degree of conservation also contains the residues that make up the dimeric interface. Up to 24 of the 27 residues within 3 Å of their dimer partner are fully conserved among the viruses that infect old world monkeys, apes, and humans. Additionally, these structure studies indicate the presence of a tetramer, which has been hypothesized to be a consequence of ICP0 dimer formation for some time^55,56^. This tetramer appears to extend the β-barrel formation and forms a hole in the middle that is approximately 11 Å x 9 Å wide and 15 Å deep. Further analyses can help to determine the nature of this interaction and its importance in viral transactivation, ubiquitination, and HSV-1 reactivation.

Small β-barrels, like the one described in this study, are a unique and highly diverse class of protein folds. Despite the major differences in sequence as a class of proteins, the process by which they fold and perform seems to be indifferent to these variations^51^. As such, small β-barrels serve a large variety of tasks, from RNA biogenesis to regulation of signaling cascades^51,57,58^. We can hypothesize two potential functions for the ICP0 C-terminal β-barrel. It could be responsible for inter-molecular ubiquitination between monomers of ICP0. Ubiquitination as a regulatory mechanism to control the amount of ICP0 present during lytic infection has been observed with the ICP0 binding protein, ubiquitin-specific protease 7 (USP7)^17^. Moreover, ICP0 can regulate USP7 through its ubiquitination and subsequent proteasomal degradation^17^. Future experiments examining ICP0’s dimerization domain in relation to self-ubiquitination will help to better understand how it regulates ICP0’s stability.

Furthermore, it is plausible that this motif may recognize certain histone tail modifications within chromatin. It has been reported that specific histone tail modifications are associated with HSV-1 reactivation and latency^59^. Chromatinization of viral DNA can occur when HSV-1 infects a cell, and specific post-translational modifications of histones can form chromatin-repressing structures to prevent viral genes from being expressed. ICP0 has been reported to interfere with this chromatinization by reducing the number of histones that associate with the viral DNA and promoting histone acetylation, linked to transcription (reviewed in ^60^).

It is likely that the extensive contact between monomers of ICP0 to form a dimer inhibits the binding of other interacting partners. The unique dimeric structure of ICP0 reveals that it likely makes a decision: either to bind to another monomer of itself or another partner such as SUMO1 or SUMOylated proteins. Of ICP0’s seven SUMO-binding domains, three are within the C-terminal domain and located in the dimeric interface (i.e., SLS5, SLS6, and SLS7). Based on computer modeling, the β-strands of ICP0 that interact with SUMO1 do so either in parallel or antiparallel to the interacting β-strand of SUMO1. According to the binding models described in this report, it appears SUMO1 binding blocks the sites that would normally interact for dimerization. It is known that SUMOylation of PML, a constituent of ND10 bodies, which is targeted for degradation by ICP0^19,22,23,61^. Interestingly, the IE protein and major transcription factor of HSV-1, ICP4, also binds in this region of ICP0^62^, but it is unclear at this point whether ICP0 dimerization affects this interaction. Collectively, these data suggest that ICP0 regulates at least two of its functions in an interdependent manner via dimerization.

Finally, this structure lies within an important multi-functional region of ICP0 (Figure 1). Consequently, it is tempting to speculate that inhibition of ICP0 dimer formation by small compounds could impair HSV-1 lytic replication. Notably, ICP0 has been used as a target of viral inhibition in at least two previous studies. A published report from our group using a high-throughput assay identified 70 potential inhibitors of ICP0’s transactivation activity, which included inhibitors of cyclin-dependent kinase (cdk) activities, trichothecenes (a secondary metabolite from fungi), and lipopeptide biosurfactants^63^. In another study using an *in vitro* high-throughput screen, 3,4,5-aryl-substituted isoxazole was identified as a potential inhibitor of ICP0’s E3 ubiquitin ligase activity^64^. Future experiments will assess the possibility of identifying chemical probes that disrupt the dimer interface of ICP0.

## CONCLUSION

In conclusion, we have characterized the first large structure of the HSV-1 ICP0 C-terminal dimer domain. The dimer structure, which contains two small β-barrels, represents a novel fold, and supports previous studies suggesting that ICP0 can also form a multimeric complex. Structural modeling indicates that due to steric hindrance, ICP0 must decide to either dimerize or to bind interacting partners such as SUMO1. While the exact function of dimerization remains elusive, this crystal structure will assist us in examining the role that dimer formation plays in ICP0 activities during HSV-1 replication and reactivation.

## STRUCTURE DEPOSITION

The coordinates and structure factors for ICP0 HE and His-ICP0 have been uploaded to the Worldwide Protein Databank (wwPDB) with the accession codes 8UA2 and 8UA5, respectively.

## DATA AVAILABILITY STATEMENT

The data for this study are available from the corresponding author upon suitable request.

## AUTHOR CONTRIBUTIONS

**Erick McCloskey:** investigation; writing-original draft; visualization; writing-review and editing; data curation. **Maithri Kashipathy:** investigation; methodology. **Anne Cooper:** Investigation; writing-review and editing; formal analysis. **Philip Gao:** Investigation; writing-review and editing; conceptualization; formal analysis; resources; supervision; data curation. **David K. Johnson:** Investigation; methodology; validation; visualization; writing-review and editing; data acquisition; formal analysis; resources; conceptualization; data curation. **Kevin P. Battaile:** Investigation; data acquisition; resources. **Scott Lovell:** Investigation; methodology; validation; visualization; writing-review and editing; data acquisition; formal analysis; supervision; resources; conceptualization; data curation. **David J. Davido**: Conceptualization; investigation; funding acquisition; writing-original draft; writing-review and editing; project administration; supervision; resources; visualization; validation; data curation.

## ACKNOWLEDGEMENTS

This work was supported by the University of Kansas, NIH Graduate Training at the Biology-Chemistry Interface Grant T32 GM132061 from the NIGMS, COBRE-PSF NIH P30 GM1110761, and COBRE-CBID NIH P20 GM113117. Use of the IMCA-CAT beamline 17-ID at the Advanced Photon Source was supported by the companies of the Industrial Macromolecular Crystallography Association through a contract with Hauptman-Woodward Medical Research Institute. Use of the Advanced Photon Source was supported by the U.S. Department of Energy, Office of Science, Office of Basic Energy Sciences under contract no. DE-AC02-06CH11357.

