## Supplemental Data for "HSV-1 ICP0 Dimer Domain Adopts a Novel β-barrel Fold"

| | ICP0 LE | ICP0 HE | His-ICP0 |
|---|---|---|---|
| **Data Collection** | | | |
| Unit-cell parameters (Å, °) | *a*=*b*=96.60, *c*=75.33 | *a*=*b*=95.91, *c*=74.53 | *a*=*b*=95.15, *c*=76.72 |
| Space group | $P4_12_12$ | $P4_12_12$ | $P4_12_12$ |
| Resolution (Å)[†] | 46.36-3.00 (3.18-3.00) | 47.95-2.65 (2.78-2.65) | 43.45-2.45 (2.55-2.45) |
| Wavelength (Å) | 1.5895 | 1.0000 | 1.0000 |
| Temperature (K) | 100 | 100 | 100 |
| Observed reflections | 171,401 | 134,156 | 89,786 |
| Unique reflections | 7,423 | 10,603 | 14,026 |
| <I/(σI)>[†] | 13.1 (1.7) | 14.6 (1.9) | 13.1 (1.6) |
| Completeness (%)[†] | 98.0 (100) | 100 (99.9) | 99.9 (100) |
| Multiplicity[†] | 23.1 (24.9) | 12.7 (13.4) | 6.4 (6.3) |
| $R_{merge}$ (%)[†, ‡] | 27.6 (263.8) | 12.3 (145.2) | 11.3 (92.6) |
| $R_{meas}$ (%)[†, ¶] | 28.2 (269.3) | 12.8 (150.7) | 12.4 (101.2) |
| $R_{pim}$ (%)[†, ¶] | 5.8 (53.6) | 3.6 (41.0) | 4.8 (40.2) |
| $CC_{1/2}$ [†, ‖] | 0.997 (0.749) | 0.999 (0.862) | 0.998 (0.714) |
| DelAnom CC[#] | 0.333 | 0.067 | |
| **Refinement** | | | |
| Resolution (Å)[†] | | 34.73-2.65 | 43.45-2.45 |
| Reflections (working/test)[†] | | 10,033/511 | 13,323/667 |
| $R_{factor}$ / $R_{free}$ (%)[†,§] | | 21.8/28.8 | 19.2/25.4 |
| No. of atoms (Protein/Iodide/water) | | 1,684/2/- | 1,743/3/42 |
| **Model Quality** | | | |
| R.m.s deviations | | | |
| Bond lengths (Å) | | 0.009 | 0.009 |
| Bond angles (°) | | 1.136 | 0.991 |
| Average *B*-factor ($Å^2$) | | | |
| All Atoms | | 71.2 | 44.8 |
| Protein | | 71.2 | 44.8 |
| Iodide | | 106.42 | 58.5 |
| Water | | - | 42.2 |
| Coordinate error (maximum likelihood) (Å) | | 0.35 | 0.34 |

| Ramachandran Plot | | |
|---|---|---|
| Most favored (%) | 97.7 | 97.8 |
| Additionally allowed (%) | 2.3 | 2.2 |

**Supplemental Table 1:** Crystallographic data for ICP0.

† Values in parenthesis are for the highest resolution shell.

‡ $R_{merge} = {}_{hkli}\, |I_i(hkl) - \langle I(hkl)\rangle| \,/\, {}_{hkli}\, I_i(hkl)$, where $I_i(hkl)$ is the intensity measured for the *i*th reflection and $\langle I(hkl)\rangle$ is the average intensity of all reflections with indices hkl.

§ $R_{factor} = {}_{hkl}\, ||F_{obs}\,(hkl)| - |F_{calc}\,(hkl)|| \,/\, {}_{hkl}\, |F_{obs}\,(hkl)|$; Rfree is calculated in an identical manner using 5% of randomly selected reflections that were not included in the refinement.

¶ $R_{meas}$ = redundancy-independent (multiplicity-weighted) $R_{merge}$[1,2]. $R_{pim}$ = precision-indicating (multiplicity-weighted) $R_{merge}$[3,4].

|| $CC_{1/2}$ is the correlation coefficient of the mean intensities between two random half-sets of data[5,6].

### DelAnom CC is the correlation coefficient between the Bijvoet differences ($I_{(hkl)} - I_{(-h\text{-}k\text{-}l)}$) from two random half-sets of data[1] and is used to estimate the anomalous signal strength.

| β-strand group | Residue N-Atom | Residue O-Atom | Distance (Å) |
|---|---|---|---|
| βG1 | Y643 | L747 | 2.96 |
| | L644 | V653 | 2.91 |
| | I646 | S651 | 3.01 |
| | L653 | L644 | 2.85 |
| | M734 | G738 | 2.83 |
| | G738 | M734 | 2.69 |
| | M740 | L732 | 2.84 |
| | L741 | V748 | 2.76 |
| | D743 | T746 | 2.79 |
| | T746 | D743 | 2.82 |
| | V748 | L741 | 2.95 |
| βG2 | D666 | L684 | 2.98 |
| | L668 | V682 | 2.77 |
| | I670 | A680 | 2.92 |
| | D672 | N677 | 3.06 |
| | G676 | D672 | 2.84 |
| | V682 | L668 | 2.84 |
| | L684 | D666 | 2.88 |
| βG3 | H715 | R759 | 2.99 |
| | T717 | R757 | 2.74 |
| | R759 | H715 | 2.87 |

**Supplemental Table 2:** Backbone hydrogen bond interactions in the β-strand regions of the ICP0 C-terminal dimer domain.

| Subunit: AB Residue/Atom | Subunit: CD Residue/Atom | Distance (Å) |
|---|---|---|
| A: S731/N | D: N739/OD1 | 2.8 |
| A: W733/N | D: M740/O | 2.84 |
| A: G745/N | D: S650/OG | 2.68 |
| B: T735/N | C: N730/OD1 | 2.98 |
| B: M740/N | C: S731/O | 3.08 |
| B: F742/N | C: W733/O | 3.01 |
| B: Q744/N | D: E674/OE2 | 2.87 |
| B: N739/OD1 | C: S739/N | 2.8 |
| B: M740/O | C: W733/N | 2.84 |
| B: S650/OG | C: G745/N | 2.68 |
| A: N730/OD1 | D: T735/N | 2.98 |
| A: S731/O | D: M740/N | 3.08 |
| A: W733/O | D: F742/N | 3.01 |
| B: E764/OE2 | D: Q744/N | 2.87 |

**Supplemental Table 3:** Hydrogen bond interactions between subunits in the ICP0 tetramer (dimer of dimers). Highlighted interactions occur between the β6-β7 strands.

**A**

**B**

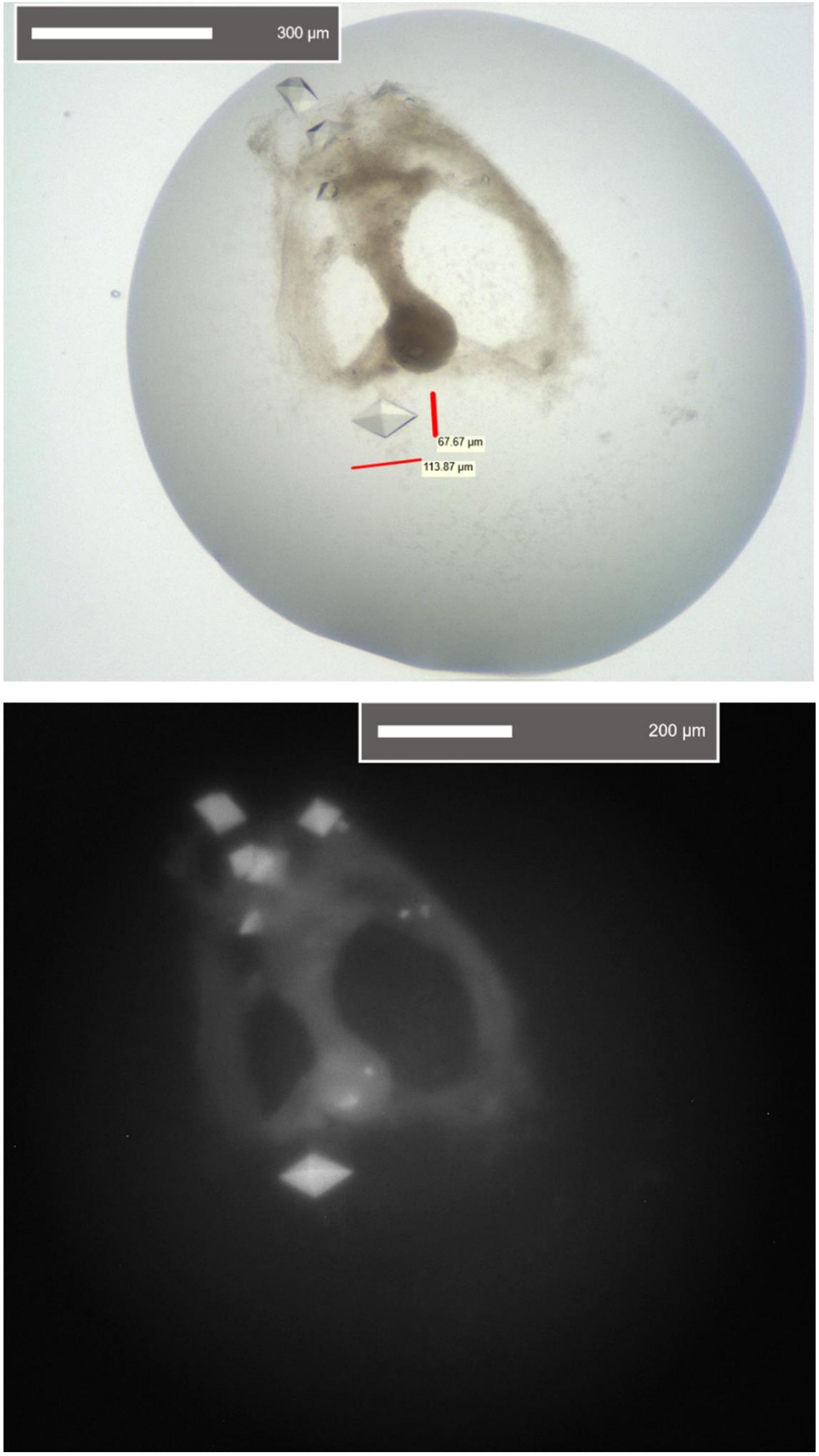


**Supplemental Figure 1:** Crystals of ICP0. **A)** Visible light image and **B)** UV fluorescence image.

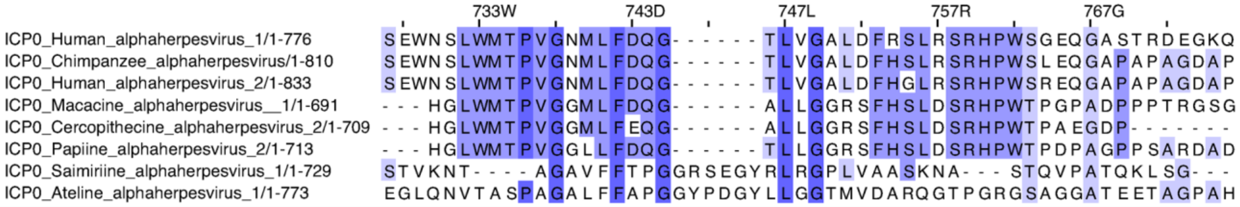

**Supplemental Figure 2:** Alignment of ICP0 CTD tetramer interface residues of unique herpesvirus species containing the CTD. The numbering is relative to the HSV-1 ICP0.

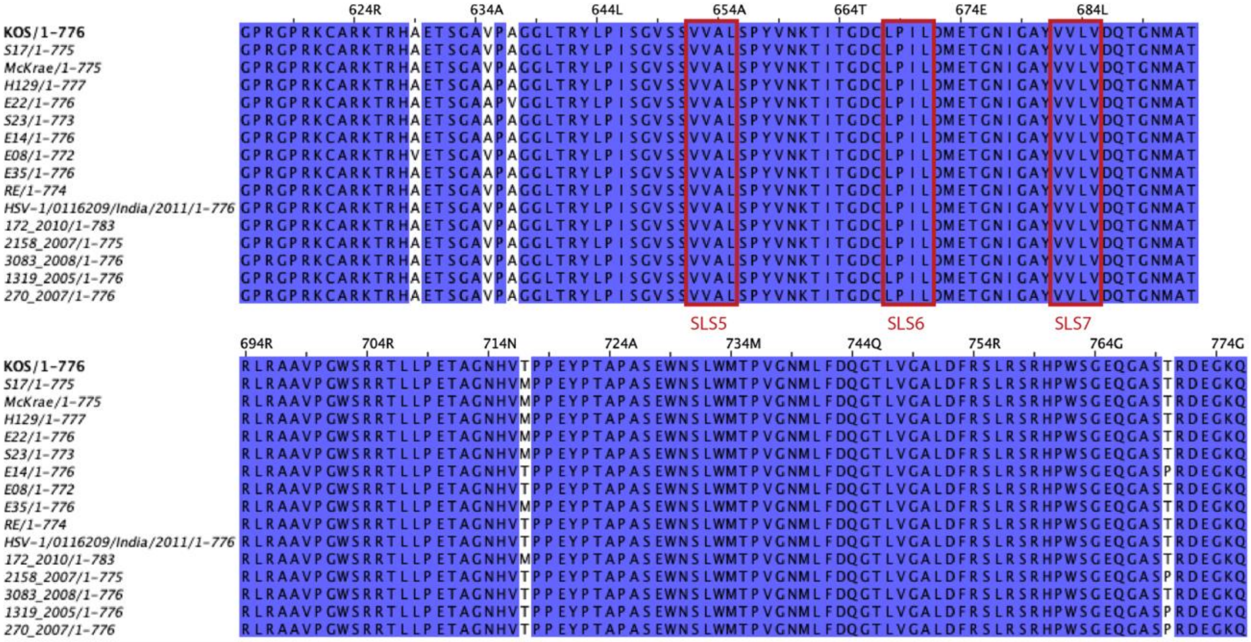


**Supplemental Figure 3:** Alignment of full ICP0 CTD of HSV-1 strains and clinical isolates. The full ICP0 CTD of various HSV-1 strains and clinical isolates. The numbering is relative to the HSV-1 ICP0 KOS strain.

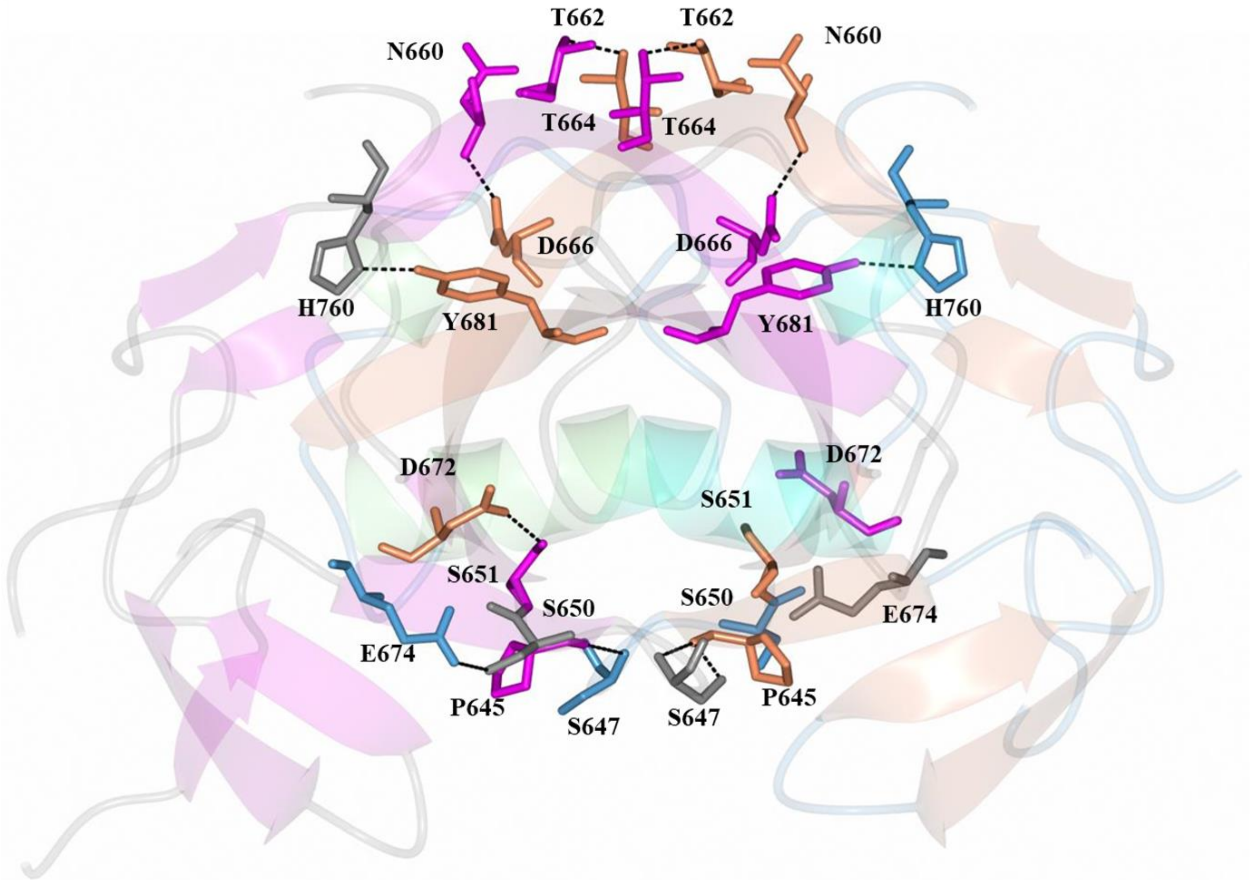


**Supplemental Figure 4:** Side chain hydrogen bond interaction between subunits (dashed lines). Subunit A helices (cyan), β-strands (magenta) and loops (gray). Subunit B helices (green), β-strands (tan) and loops (blue).

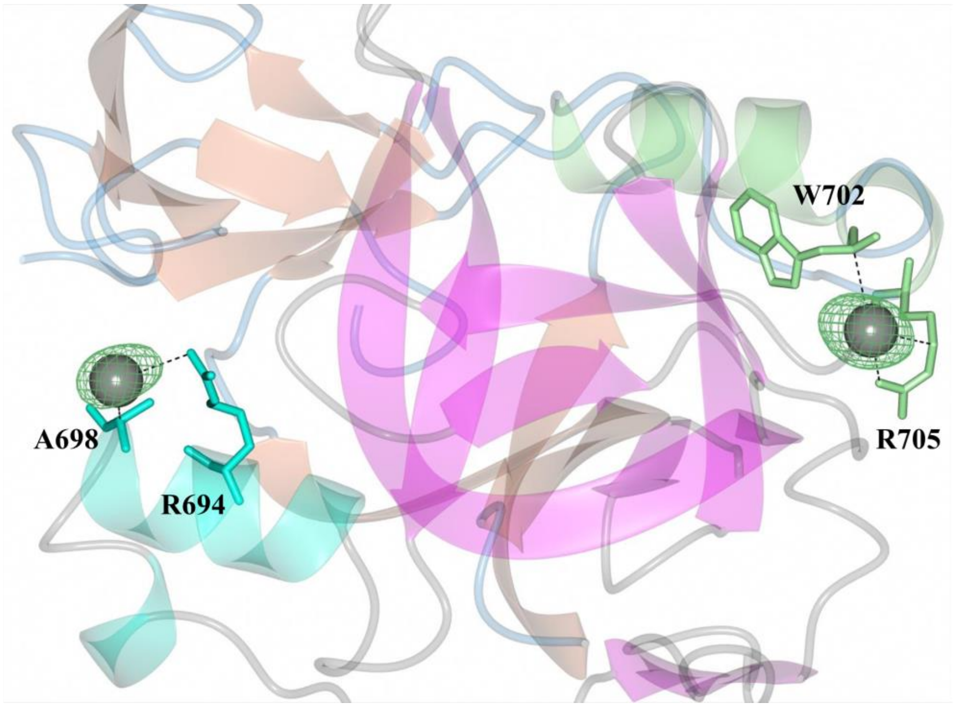


**Supplemental Figure 5:** Phased anomalous difference map (green mesh) contoured at 3 showing the positions of the iodide ions (gray spheres). The dashed lines indicated close contacts between 3.6-3.9 Å.

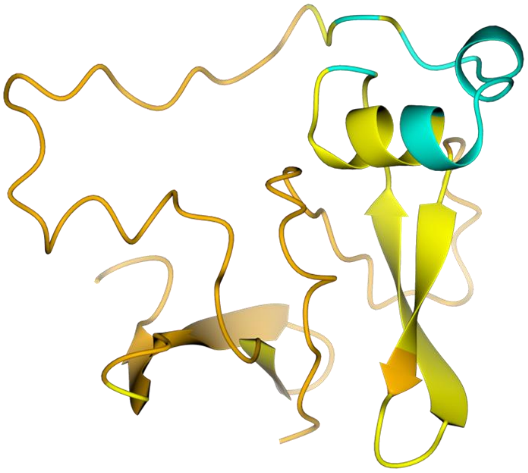

**Supplemental Figure 6:** Alphafold model of the monomeric ICP0 CTD. The Alphafold model of the ICP0 CTD is shown in ribbons, colored by the pLDDT (confidence): confident (cyan, 90 > pLDDT > 70), low (yellow, 70 > pLDDT > 50), and very low (orange, pLDDT < 50).

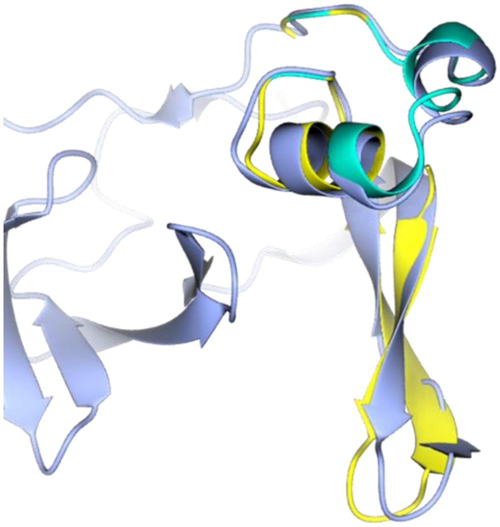

**Supplemental Figure 7:** The region of the Alphafold model of the monomeric ICP0 CTD with low or moderate confidence recapitulates the monomeric structure. The Alphafold model of the ICP0 CTD with pLDDT > 50 is shown in ribbons (cyan, 90 > pLDDT > 70 vs yellow, 70 > pLDDT > 50). A single chain of the ICP0 CTD is shown in silver.

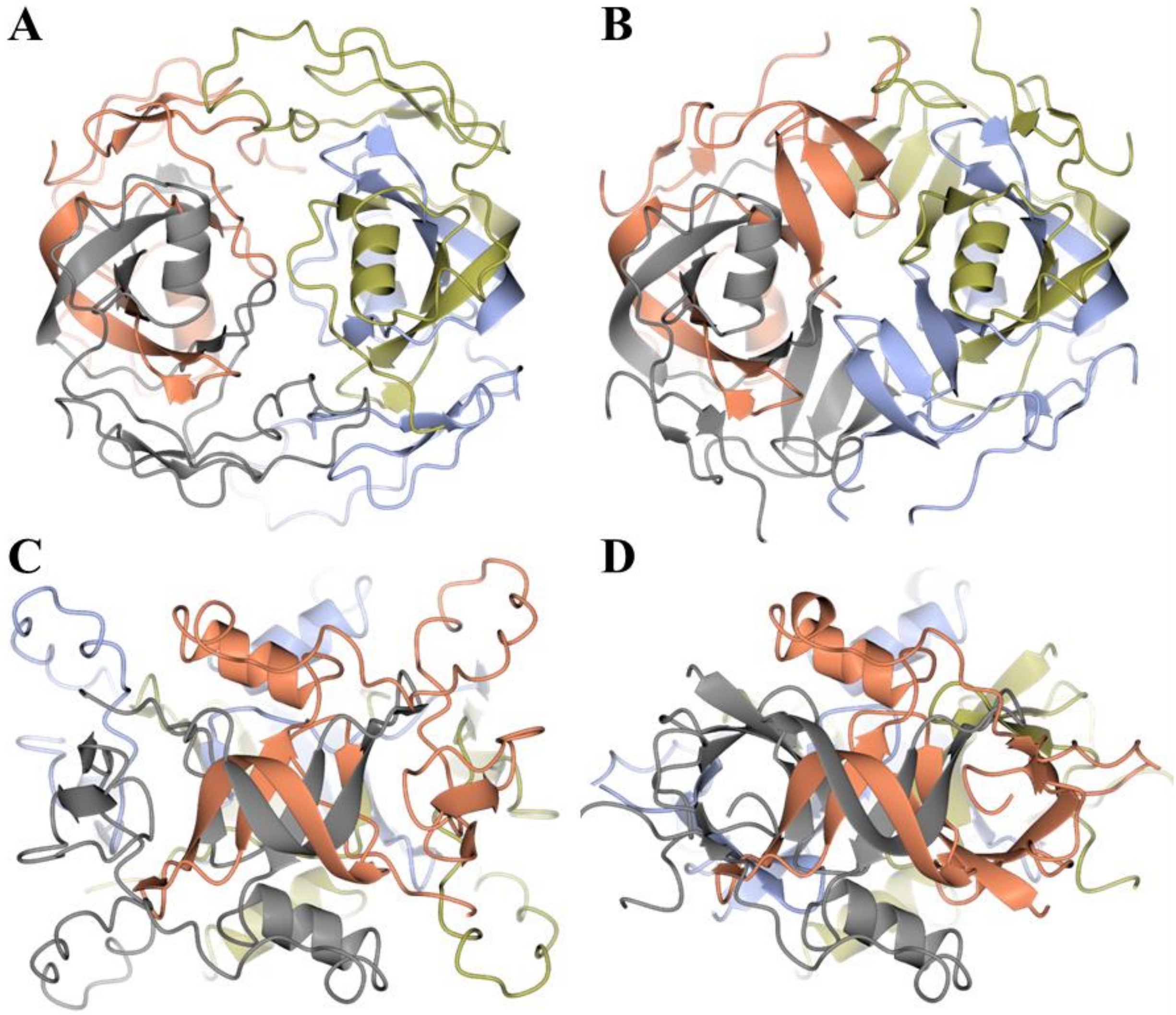


**Supplemental Figure 8:** Alphafold model of the tetrameric ICP0 CTD compared to the solved structure. The Alphafold model of the ICP0 CTD tetramer is shown in ribbons on the left (**A and C**), while the crystal structure is shown on the right (**B and D**), colored by the chain. The dimeric interface, particularly the twisted β-strands, were modeled accurately (**A and B**). However, generally none of the strands comprising the stacked barrels were modeled (**C and D**).

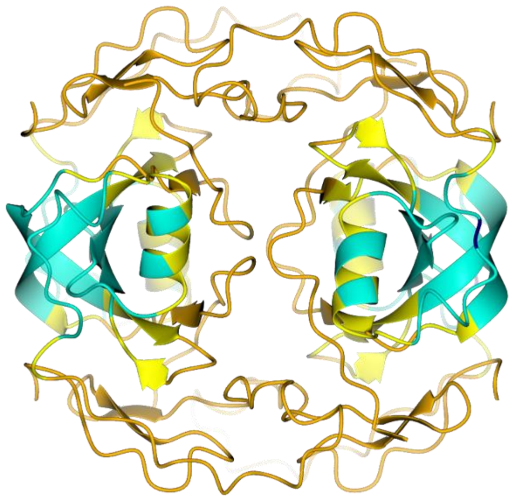

**Supplemental Figure 9:** Confidence of the Alphafold model of the tetrameric ICP0 CTD. The Alphafold model of the ICP0 CTD is shown in ribbons, colored by the pLDDT (confidence): confident (cyan, 90 > pLDDT > 70), low (yellow, 70 > pLDDT > 50), and very low (orange, pLDDT < 50).

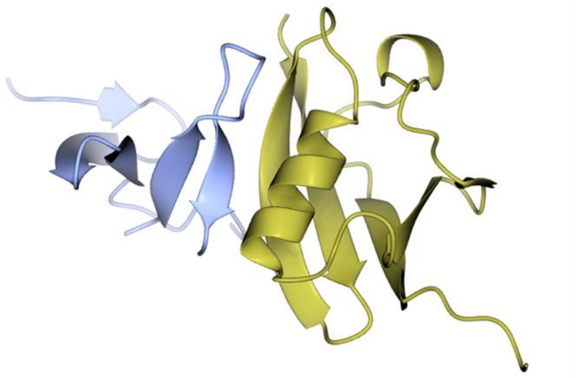

**Supplemental Figure 10:** A model of SUMO binding anti-parallel to ICP0 at SLS5. A folded subdomain containing SLS5 is shown in silver ribbons, while SUMO is represented by gold ribbons.

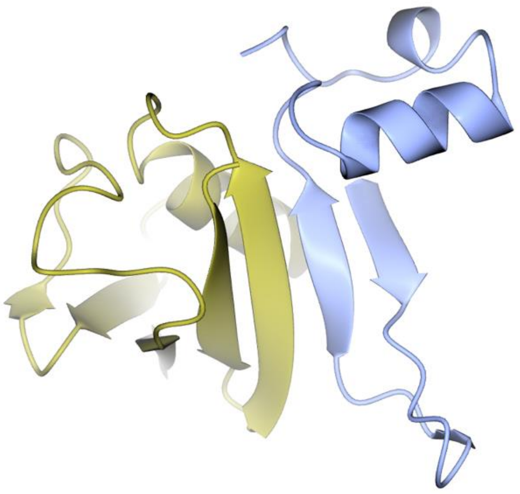

**Supplemental Figure 11:** A model of SUMO binding parallel to ICP0 at SLS7. A folded subdomain containing SLS7 is shown in silver ribbons, while SUMO is represented by gold ribbons.
